# Developing critical thinking in STEM education through inquiry-based writing in the laboratory classroom

**DOI:** 10.1101/686022

**Authors:** Ah-Jung Jeon, David Kellogg, Mohammed Asif Khan, Greg Tucker-Kellogg

**Author notes:** Correspondence: Greg Tucker-Kellogg < >.

## Abstract

Laboratory pedagogy is moving away from step-by-step instructions and toward inquiry-based learning (IBL), but only now developing methods for integrating inquiry-based writing (IBW) practices into the laboratory course. Based on an earlier proposal (Science 332:919 (**2011**)), we designed and implemented an IBW sequence in a university bioinformatics course.

We automatically generated unique, double-blinded, biologically plausible DNA sequences for each student. After guided instruction, students investigated sequences independently and responded through IBW writing assignments. IBW assignments were structured as condensed versions of a scientific research paper, and because the sequences were double blinded, they were also assessed as authentic science and evaluated on clarity and persuasiveness.

We piloted the approach in a seven-day workshop (35 students) at Perdana University Graduate School of Medicine in Kuala Lumpur. We observed dramatically improved student engagement and indirect evidence of improved learning outcomes over a similar workshop without IBW. Based on student feedback, initial discomfort with the writing component abated in favor of an overall positive response and increasing comfort with the high demands of student writing. Similarly encouraging results were found in a semester length undergraduate module at the National University of Singapore (155 students).

## Introduction

The dominant teaching model for post-secondary Science, Technology, Engineering, and Mathematics (STEM) education pairs classroom lectures with regular laboratory sections (Beichner, 2014; Lunetta, Hofstein, & Clough, 2007). In this model, lectures provide most of the course content, while the accompanying laboratories provide an opportunity for students to solve problems, gain skills, and reinforce what they have learned in the lectures. What counts as a laboratory in this scenario varies widely, but typically the work of the laboratory classroom is meant to mimic the work of the science under study. Because laboratory sections are often restricted in size — either due to resource limitations or to enable group work, discussion, and reflective activity — several lab sections may be needed to pair with a single large lecture. Lectures and lab are often assessed separately, with lectures assessed through graded exams or homework, and labs assessed through evaluation of written laboratory reports. Classroom assignments, laboratory exercises, and laboratory reports share the objectives of providing students with scientific education and training by exposing them to simulated, simplified, and well-controlled scientific problem situations.

Lecture-based pedagogy is in the midst of a decades-long effort to adopt more active learning, in which students construct their own understanding in the course of learning activities instead of receiving the learned perspective of the instructor. The term *active learning* is an umbrella concept comprising a range of alternatives to the expository lecture, including but not limited to research-based, problem-based, and inquiry-based learning. A 1999 review offers a still useful taxonomy of these categories, distinguishing among laboratory instruction styles by outcome, approach, and procedure (Domin, 1999). Research-based learning engages students in active research problems. Research-based laboratory classrooms, encouraged by the AAAS report *Vision and Change in Undergraduate Biology Education*, increase student engagement and retain the uncertainty of authentic scientific inquiry (AAAS, 2011). Problem-based learning, more often implemented in introductory courses, works in a more constrained problem context than research-based laboratories. Inquiry-based learning (IBL) reflects the pedagogical understanding that authentic scientific activity arises out of an ongoing practice of inquiry, and that modeling inquiry, including the formation of appropriate questions as well as of sound and reasonable answers, is one way to model authentic science. While the cookbook-style classroom lab presumes both the design and the desired outcome of the experimental scene, IBL classrooms provide alternative scenarios of varying degrees of openness (Anderson, 2007).

Data to support the turn toward active learning within post-secondary education is substantial: a recent meta-analysis of 225 studies has shown that active learning in undergraduate science classes increases student performance when compared to expository lecture-based learning (Freeman et al., 2014). Active learning has retained its appeal in a period of rapid technological change. The dissemination of personal technology, for example, while it may have problematic effects on attention and behavior (see Reid, 2018), can both challenge the traditional authority of the lecture and provide new opportunities for active interaction (Carloye, 2017). By the same token, at least under carefully monitored conditions, hybrid or online instruction may be seen as facilitating active learning rather than hindering it (Baepler, Walker, & Driessen, 2014).

Compared to the science classroom lecture, learning activities in the laboratory have been slow to change despite evidence that reform to laboratory pedagogy can be effective (Weaver, Russell, & Wink, 2008). This is not to say the classroom laboratory has been unaffected by reform; indeed, the term *cookbook* is now applied almost entirely as a pejorative for unreformed traditional laboratory instruction. Nevertheless, reform in laboratory learning activities has been uneven (Beck, Butler, & Burke da Silva, 2014). There are obvious practical challenges — such as cost and safety issues — to reforming laboratory-based learning activities. Beyond practical matters, activities in the laboratory may be slow to change because they already have the appearance of active learning. What could be more active than a group of students beavering away in the laboratory and writing reports on their work? Isn’t laboratory learning *always* and *already* active learning? Despite this appearance, the traditional cookbook lab has been challenged by research-based, problem-based, and inquiry-based laboratory reforms.

Students in classroom laboratories are commonly assessed through the writing they produce. In the traditional lab report, students describe objectives, methods, and results, and sometimes produce a discussion; this structure is meant to imitate the activities of scientists producing science. A common alternative to the traditional lab report uses the Science Writing Heuristic (SWH), where students are asked to answer specific and more focused questions that directly address the main learning objectives (Keys, Hand, Prain, & Collins, 1999). The SWH has the apparent advantage of minimizing the redundancies of some components in typical lab reports. Often, a hybrid form is adopted whereby the assignments largely take the form of a lab report but also include short answer questions in the requirement. Depending on how they are structured, both lab reports and the SWH may — or may not — be examples of writing to learn (WTL), where the activity of writing reinforces and reframes the course content for the student learner. Although WTL tends to favor low-stakes and reflective writing activities (Kalman, 2008), structured questions such as the SWH and lab reports requiring deliberation can also be used to meet WTL objectives (see Chapter 7 of Kalman, 2017). When students write about newly acquired knowledge, they may develop a deeper understanding of major concepts, especially when writing assignments require students to analyze and synthesize information. However, laboratory reports do not *necessarily* result in more engaged student learning than activities not requiring writing, nor do laboratory reports necessarily lead to more effective assessment.

The main hindrance to such engagement is the lack of authenticity: in the cookbook lab, and even in some mildly reformed labs, both students and instructors know what the lab report should “correctly” say before having to write or read one. In such cases, students may focus their work on matching the contents of their reports to the expectations of their instructors, and instructors may evaluate student reports in kind. The reports that emerge are unlikely to display either in-depth analysis or reflection on learning; when students are all performing the same task, as many experienced laboratory instructors can attest. In a laboratory classroom built around inauthentic problems, both students and instructors will tend to focus on the correctness of the answers. In an authentic inquiry-based learning environment, by contrast, there is at least the possibility that students will construct meaning for themselves, modify preexisting concepts, and enrich their understanding by engaging with context and with others (Anderson, 2007).

The primary site of reform in IBL laboratory courses, including research-based and problem-based labs, has been the laboratory activity itself, with one review finding a predominance of guided rather than open-ended inquiry activities (Beck et al., 2014). Although IBL lab scenarios may include WTL activities, the primary assessment tool — the lab report — has remained relatively unchanged. One proposal (Moskovitz & Kellogg, 2011) suggested developing inquiry-based writing (IBW) assignments that would address limitations to inquiry found in the traditional lab report. To implement IBW fully in the laboratory classroom, both the nature of the problem and the nature of of the assessment need to be changed. Students should be given authentic problems whereby no fixed answers are known to anyone, including the instructors, and no two individuals receive exactly the same problem. The resulting lab reports should also be unique to each student and blinded to teaching staff. Assessment of IBW in this implementation would use rubrics based on the written argument and evidence, much like an actual scientific manuscript review. In IBW, writing is skillful or unskillful, convincing or unconvincing, rather than closer to or farther from a closely guarded correct response.

In this paper, we report on the implementation of an inquiry-based writing sequence for university-level bioinformatics education at two sites. We generated double-blinded DNA sequence-based problems for student inquiry. Each student was provided a different sequence as a basis for inquiry and was assessed through a set of increasingly challenging individual (and later group) IBW assignments. We examineed student performance on written tasks as well as student response to the class experience (through informal surveys and structural topic modeling of student evaluations). Together, these results suggest that an IBW assignment sequence, when carefully constructed and rigorously implemented, can enrich course content and lead to a module that is both challenging and satisfying for students. We examine possibilities and difficulties for future IBW development along these lines.

## Methods

### Educational setting

Variations of inquiry-based writing laboratory discussed in this paper were implemented twice, in different locations and with different durations. The first was a week-long full time continuing education workshop organised by Perdana University in Kuala Lumpur, Malaysia. The second was a semester-long undergraduate course (module) designed for second year students majoring in Life Sciences at the National University of Singapore (NUS). We will refer to the first implementation as the *pilot* and the second implementation as the *course* (Table 1).

**Table 1:** Comparison of settings for the two implementations discussed in this work.

|  | Pilot | Course |
| --- | --- | --- |
| **Setting** | Perdana University | National University of Singapore |
| **Cohort** | Continuing education students | NUS undergraduate students |
| **Duration** | 8 days | one semester (13 weeks) |
| **Lecture hours** | 14 | 26 |
| **Laboratory hours** | 24 | 44 |
| **Class size** | 30 students | 155 students |
| **Format** | Interleaved lectures and labs each day | Separate lecture and lab sessions each week |

### Week long workshop (pilot)

The pilot was conducted at an eight day introductory bioinformatics workshop hosted by Perdana University in Kuala Lumpur, Malaysia in July 2013. The workshop was advertised as a continuing education opportunity for professionals. The backgrounds of the thirty participants varied from a recent high school graduate to researchers in related industries. Most, but not all, participants were from Malaysia. Assessment included two short individual written reports.

Each full workshop day was organised with interspersed lectures and practical laboratories. We introduced the assessment structure and the inquiry-based writing assignments on the first day of the workshop, and assigned the first inquiry-based writing assignment on day 3. Assignments were due in the morning, marked during the day, and handed back at the end of the day.

### One semester module (course)

After the pilot, we implemented IBW into LSM2241, Introductory Bioinformatics, at the National University of Singapore during the 2013/2014 academic year. LSM2241 is a semester-long (13 week) major elective for second year life sciences majors. The class included two hours of lectures and one four hour laboratory session per week. In prior iterations of the course, students were assessed through an oral team-based presentation on a topic, a midterm test, and a final exam. With the introduction of IBW, students were assessed through two individual written reports, one group written report, a midterm, and a final exam. All lectures were given by the same instructor (GTK) both during the prior semester and after the introduction of IBW.

### Contents and pedagogical structure

Both implementations were built around a theme of sequence comparison viewed through the lens of molecular evolution. Topics included literature databases and reference management software, pairwise and multiple sequence alignment, BLAST and related database search, basic molecular phylogenetics, and protein structure comparisons. Each lecture was paired with a laboratory session, some of which were inquiry-based. In conventional and problem-based laboratory sessions, students practised what they learned and explored problem sets to reinforce the content of the lectures.

While the week-long pilot and the semester-long course covered similar material, they did so at different depths and with different expectations of students. The course implementation included both more theory and more discussion of, for example, algorithmic issues than did the pilot.

### Inquiry-based writing assignments

Three IBW assignments were developed for the pilot and expanded for the course, as shown in Table 2. The IBW assignments were designed to be linked: the work of the later assignments was built on the results of the earlier assignments. This linkage was intended to allow the students to use their inquiry-based sequences throughout the semester.

**Table 2:**
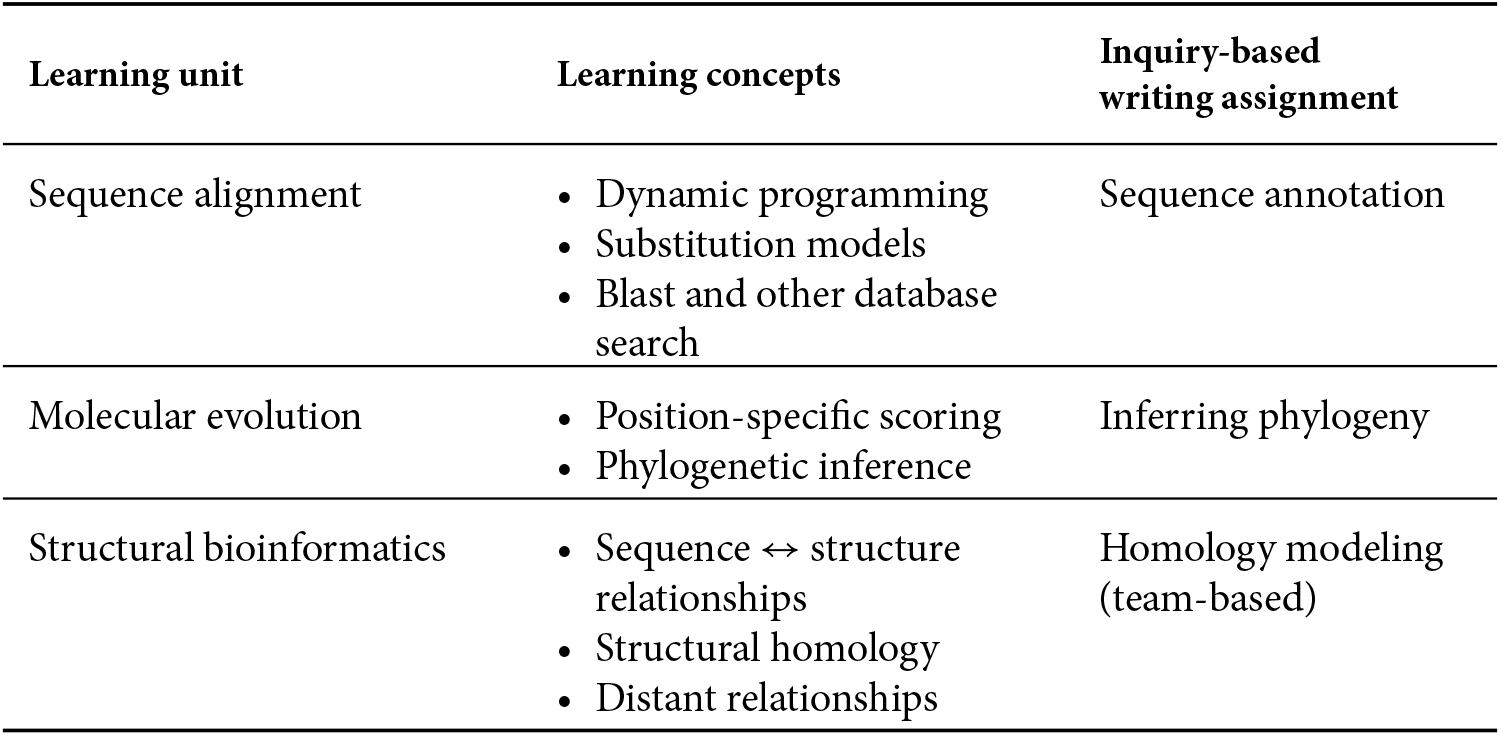
Learning units of LSM2241, Introductory Bioinformatics, and inquiry-based writing assignments for each unit.

| Learning unit | Learning concepts | Inquiry-based writing assignment |
| --- | --- | --- |
| Sequence alignment | • Dynamic programming<br>• Substitution models<br>• Blast and other database search | Sequence annotation |
| Molecular evolution | • Position-specific scoring<br>• Phylogenetic inference | Inferring phylogeny |
| Structural bioinformatics | • Sequence ↔ structure relationships<br>• Structural homology<br>• Distant relationships | Homology modeling (team-based) |

For inquiry-based problems, individual students were provided unique but biologically plausible DNA sequences whose identities were unknown to both students and the teaching staff. During the inquiry-based portion of the practicals, students were expected to come up with their own inquiries and design how to employ the relevant bioinformatics tools to answer their research questions.

After each learning unit (Table 2), students submitted a written report in a scientific report format to present and discuss their findings. The assignments were not standalone pieces of work, but could be built on from the previous assignments. The third (team-based) required students to reflect as a team on their previous findings and write about their learning.

### Assessment of inquiry-based writing reports

The IBW assignments were assessed using rubrics designed specifically for each assignment, and students were introduced to the rubrics in advance of each assignment (course assignments can be found in the Supplementary materials). Each rubric provided expectations with examples of “Excellence” (the highest possible mark) for each criterion. All assignments included assessment criteria for rhetorical expectations, writing clarity, and proper use of literature references. Students were expected to use some form of reference management software, as covered in the first (conventional) laboratory session. The marking rubric for the group assignment included a teamwork component and a reflection component.

Students were provided a maximum page limit and twelve point font size requirement, rather than a maximum word count, for each assignment. Figures, tables, references, and supplemental data were not counted against this limit.

Teaching assistants were first trained for assignment marking by GTK and AJJ. A randomly selected subset of student submissions was reviewed by all teaching staff before marking the entire cohort for each assignment. For the pilot, IBW assignments were marked independently by the lead teaching assistant (AJJ) and the lead instructor (GTK). For the course, each assignment was marked independently by two teaching assistants, and assessments were moderated by the lead instructor (GTK). Student submissions were anonymous, identifiable only by student ID number. The assessment process was as transparent as possible. After each assignment, the graded reports were returned to students with comments provided by the teaching staff to aid student reflection, and part of one laboratory session was then devoted to review and discussion.

### Generating individual and authentic sequences for inquiry

DNA sequences for inquiry were derived using a strategy to provide unique sequences consistent with the learning objectives of the course, as outlined in Figure 1. Individualised sequences were derived from actual sequences starting from a recently published transcriptome study of the vampire bat *Desmodus rotundus* (Francischetti et al., 2013), using a collection of Perl and R scripts. DNA sequences were translated to protein and mutated using PAM matrix-directed mutation probabilities, which were converted back to DNA using vertebrate codon frequency tables as codon probabilities. Individualised sequences were designed to be biologically plausible, but not trivially recognisable. To ensure plausibility, mutated protein sequences were discarded if mutations altered key residues required for domain recognition. To ensure challenge, sequences were mutated to the extent that the closest ‘blastp’ hit shared less than 60% identity with the mutated protein sequence. Finally, DNA sequences were shifted to a random coding frame and (with 50% probability) converted to the reverse complement. Students were randomly assigned individual DNA sequences, and told that sequences should be treated as possibly incomplete cDNA sequences.

**Figure 1:**
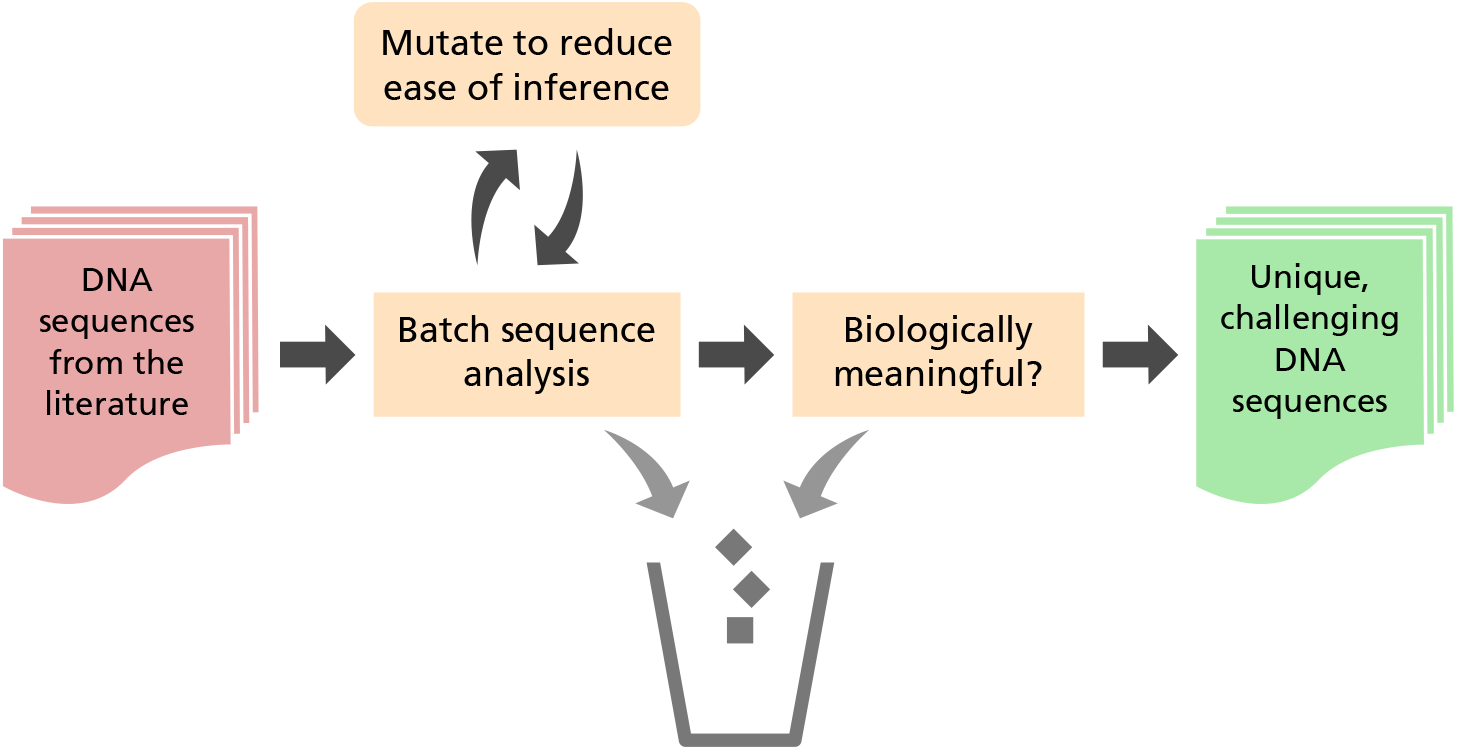
Process for random inquiry-based sequence generation

### Analysis

#### Student feedback data

Standard end-of-semester student feedback, as anonymously generated for all courses at NUS and provided to GTK as the lead instructor for these courses, included choices on a five point Likert scale, as well as free text comments on common questions. We confined our analysis of text data to student comments regarding the course, rather than comments focused on the teacher as an individual. We used limited information from these SET data, along with in-class surveys at the end of the workshop and course, to gauge student perceptions of different modes of learning. We asked students to rate both their enjoyment of the learning experience and their perception of stress on a five point Likert scale.

#### Text mining analysis of student perceptions of learning

We performed text mining of student SET comments using the stm package (Roberts, Stewart, & Tingley, 2018) in R. After stemming and removing trivial comments, we were left with 422 student comments for structural topic modeling. Each comment was labeled with the corresponding pedagogy, and the stm function run on the comments with a model that included `prevalence` ~ `pedagogy`. Plots were generated using functions in the stm package.

## Results

### Inquiry-based assignments

#### Assignment 1: sequence characterization

The first IBW assignment required students to use methods they had studied to annotate their assigned sequence. At the point of the assignment, students had studied pairwise alignment (including both Needleman-Wunch and Smith Waterman dynamic programming algorithms), comparison of sequences using dotplots, and the BLAST algorithm for sequence database search. They had also covered basic profile-based methods such as PSI-BLAST. In the assignment, students were asked to annotate the sequence using any tools covered up to that point. The wording of the assignment suggested identifying repetitive sequences, inferring homology to known sequences or families, and functional annotation (if possible). Because the sequences were based largely on incomplete mRNA sequences, and because sequences were unique, students could compare specific answers to see if they were correct, but could discuss their strategies of inquiry.

Students took a wide variety of approaches to the problem. We noticed, both in observing the class and while marking this first assignment, that the many students first attemped to apply every method they had learned, in the order they had learned it. For example, many students first calculated a dotplot of the sequence against itself. If some internal similarity was observable, some students then performed pairwise alignment of the similar regions using global or local alignment (or both, in that order). When it came to BLAST search, many students attempted all BLAST variations possible using a nucleotide query, in the order they had been introduced in the lectures: blastn, blastx, and tblastx.

Attempting the inquiry in the order concepts were covered was usually unproductive, while more productive strategies were suggested by the assignment itself. The first sentence of the assignment read “You have each been given a DNA sequence which may code for a protein, in whole or in part, in any frame.” Students who used that information to plan their inquiry often chose to start their analysis using blastx to search a database of protein sequences. For most students this was the most productive search strategy, especially when searching a well-annotated database. In contrast, most blastn and tblastx searches were uninformative, as were tools like dotplots.

The chronological order of inquiry was often reflected in a chronological organisation of the IBW report. Many students reported, and wrote about, everything they attempted, whether or not it was productive, and whether or not it served a narrative purpose. Because figures and tables were not counted against the page limit of the assignment, students often attached screenshots of all their analyses, with pages often far exceeding the main text. We interpreted that pattern among students as both a sign of discomfort with the open-ended nature of the inquiry-based assignment and unfamiliarity with different forms of scientific writing.

To address common mistakes as well as good practices, we devoted part of a laboratory session after returning the marked assignments to a focused discussion. We included isses of scientific writing in general and the differences between a laboratory notebook and a written report of discovery, as well as extensive discussions of a working hypothesis.

#### Assignment 2: phylogenetics

In the second assignment, students were required to collect a set of related sequences in order to construct a phylogenetic tree that included their assigned sequence. At the time of this assignment, the class had covered multiple sequence alignment and phylogenetic tree construction. Major student inquiry-based decisions in this assignment included what kind of tree to attempt (e.g., whether try to create a tree of a gene family or attempt to collect putative orthologs and create a species tree), what inference methods to use, how to assess tree reliability, and whether to attempt to root the tree. Depending on their choices, the discussion section of inquiry varied widely.

Again, students took a range of approaches to this problem, and some of the IBW reports uncovered student misconceptions on important topics such as homology. For example, some students argued on the basis of the basis of their results that their assigned sequence was a *particular* gene in a *particular* species, rather than bore some relation by homology. We used the review session after assignments were returned to review common mistakes and misconceptions.

One of the more interesting student findings arose from a flaw in our computational sequence generation process. Because of the dissimilarity we required in order to make the sequence annotation interesting, many students found that their assigned sequence was assigned to an especially long branch in a tree of inferred orthologs. While the origin of the sequences was unknown to the students, some students correctly inferred from this unusual tree structure that their sequence was computationally generated.

#### Assignment 3: homology modeling

The last portion of the module covered sequence-structure relationships, and included homology modelling. We had determined beforehand that a substantial minority of the assigned sequences could be used for homology modelling by students running SWISS-MODEL or MODELLER (Biasini et al., 2014; Webb & Sali, 2016) and that this could be a good basis for IBW. Unlike the first two assignments, however, we could not safely assume that all of the students had the oppotunity for a productive inquiry using their assigned sequence. Assignment 3 was therefore assigned to groups of up to five students, with the high likelihood that some of the sequences would yield interesting results.

The group setting of assignment 3 also presented an opportunity for group reflection, so we included a requirement for critical reflection on their first two assignments in the third assignment. We devoted part of one laboratory section to facilitate disussion within each group about the different approaches students had taken for their individual first two assignments.

### Student feedback on experience

We used data the standard end-of-semester student evaluations of teaching (SETs) to gauge overall views of the module and levels of difficulty. We also conducted informal, anonymous, post-course surveys of students to evaluate their experience of the laboratory. Quantitative responses included Likert-scale reports of enjoyment and stress, as well as whether students would recommend continuing the inquiry-based writing in future years. Qualitative responses included student perceptions of differences between the inquiry-based writing and conventional laboratory sessions.

From the standard SET ratings of the module (Figure 2), we noticed that introduction of inquiry-based activities was associated with improvement in student overall opinions of the module, even though student perceptions of difficulty were unchanged.

**Figure 2:**
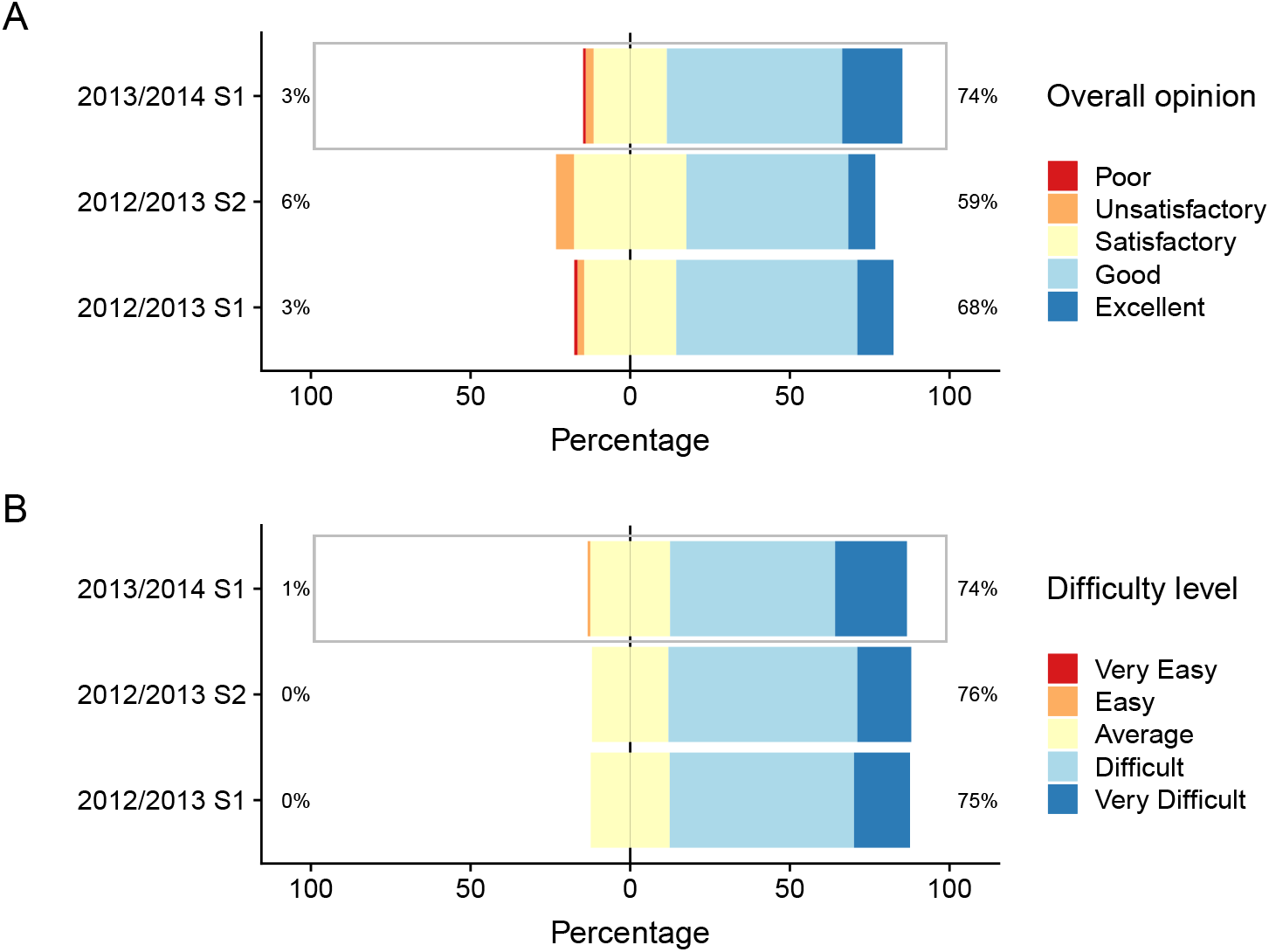
Standard responses from student evaluations of teaching. **A.** Overall opinion of the module. **B.** Perceived difficulty of the module. Responses are on a 5-level Likert scale; data are presented as stacked bar charts centered on the third level of response. The semester using inquiry-based writing is highlighted.

### The change in pedagogy was recognisable from text mining of student feedback

In addition to informal surveys, we utilised end of semester student evaluations of teaching for LSM2241. Because students could provide feedback on anything related to the course, we decided to use text mining of student feedback to see if the change in pedagogy was recognisable. For this we turned to structural topic modeling.

Topic modeling treats a collection of documents as arising from a mixture of abstract “topics”. Widely used topic modeling algorithms such as latent Dirichlet allocation (LDA) discover topics from such a document collection. We were interested in identifying topics associated with changes in pedagogy, but conventional topic modeling does not allow simultaneous discovery of topics and associations with metadata. We therefore used structural topic modeling (STM), a recent development in text mining (Roberts, Stewart, Airoldi, et al., 2014; Roberts, Stewart, Tingley, et al., 2014) designed for exactly this purpose.

We treated each non-trivial feedback comment as a document, and labeled each document according to the pedagogy used (“traditional” or “inquiry-based”). We developed a model of ten topics associated with pedagogy from 412 comments, as summarised in Figure 4A. When we examined relative topic prevalence for the ten topics, we noted that three topics were more prevalent in the IBW pedagogy, and while other topics were more prevalent in the traditional pedagogy (Figure 4B). The three topics most associated with inquiry-based learning were also highly correlated with each other (Figure 4C).

Sample comments from topic 7, the topic most associated with traditional inquiry (Figure 4D, top) show students struggling with the conceptual difficulties of the course, a repeated theme that motivated the introduction of inquiry-based writing in the first place.

In contrast, sample comments from topic 5, strongly associated with the IBW pedagogy (Figure 4D, bottom), show students commenting on the inquiry-based learning itself. Many student commented positively, or noted that it was difficult but valuable for their learning. More broadly, the topics most associated with the IBW pedagogy were filled with reflections from students on their own learning, or on the connection between the course and their experience outside of it. These topics also included complaints or negative feedback, with a common concern that assignment was unfair or varied too much between teaching assistants.

### Most students prefer inquiry-based learning, even knowing the challenge

We asked students in both the workshop and the course whether they would prefer assessment from inquiry-based writing assignments or examinations. Students responded in favour of IBW in both the workshop (29/30) and the course (48/16). For the course, we also asked students if we should continue the inquiry-based writing, and the response was overwhelmingly positive (61/72 yes, 4/72 no). Students were often very candid about their reasoning, telling us things like “even though it was difficult, it really helped us to discover the various tools used in bioinformatics”, and “It was a good learning experience which improved my writing skills.” Students who recommended against continuing IBW generally found it too difficult, or being “thrown in the deep end of the pool”.

## Discussion

### Observations of student engagement

For this paper, we implemented and evaluated a pedagogy including inquiry-based writing (IBW) in both a pilot workshop and a semester-long bioinformatics course. The appeal of IBW was motivated in part by an observed disengagement among students in previous years, in both the pilot and the course. We therefore paid attention to any anecdotal signs of change. Informal observations of the lab classroom during the inquiry-based sessions were encouraging, as it appeared that student activity was much more focused and more animated. The most discouraging behaviours from previous years, such as students using social media during class, virtually disappeared. In contrast to previous semesters, students in discussions — both with each other and with the instructors — were largely grappling with the inquiry-based problem. The students knew that the teaching staff did not know the “correct” answer to their inquiry, so the nature of the interaction was qualitatively different than in other laboratory activities.

We also noticed an elevated level of stress among students. During the pilot, which compressed considerable work into a few days, some students found the written assignments so stressful that they began to register complaints to the workshop coodinator (MAK). In response to these complaints, we decided to make the second project optional, though we offered to provide feedback and “mark” the second project for any students who turned it in. We also allowed groups to choose a traditional exercise (based on the previous year’s workshop) instead of carrying out the group-based inquiry-based assignment.

To our surprise, most students turned in and requested feedback on the optional individual inquiry, and *every* group chose to continue with the inquiry for their group project. This apparent combination of high stress and high student engagement was consistent with survey feedback in which students recorded high levels of stress despite overwhelming student preference for inquiry-based writing over conventional lab activities (Figure 3).

**Figure 3:**
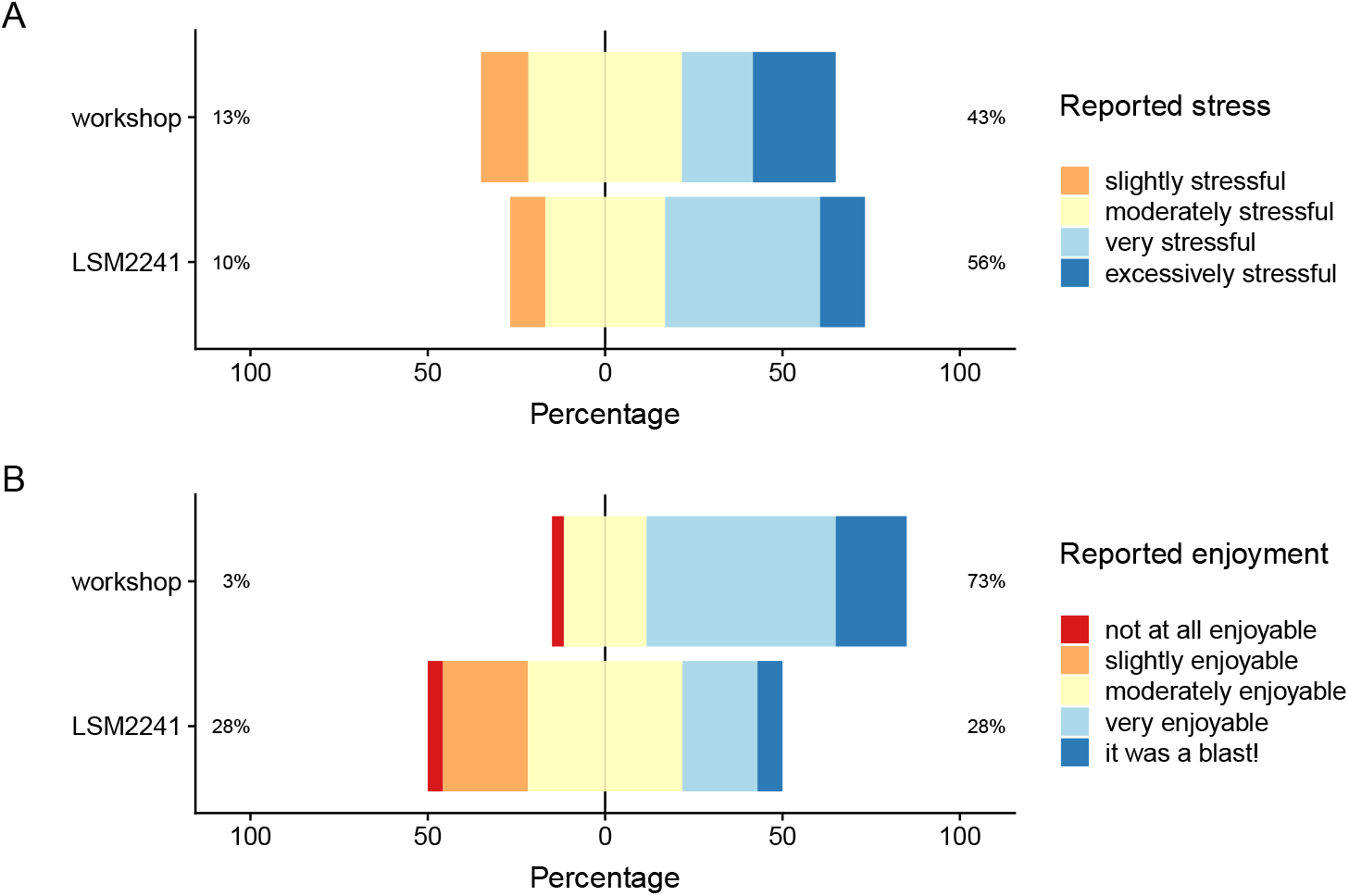
Student reports of enjoyment and stress for the workshop and the semester-long course **A** Reported stress in end of course feedback. **B** Reported enjoyment in end-of-course feedback. All stacked bar charts are centered at the third of five choices, but there were no reports of “not at all stressful”.

**Figure 4:**
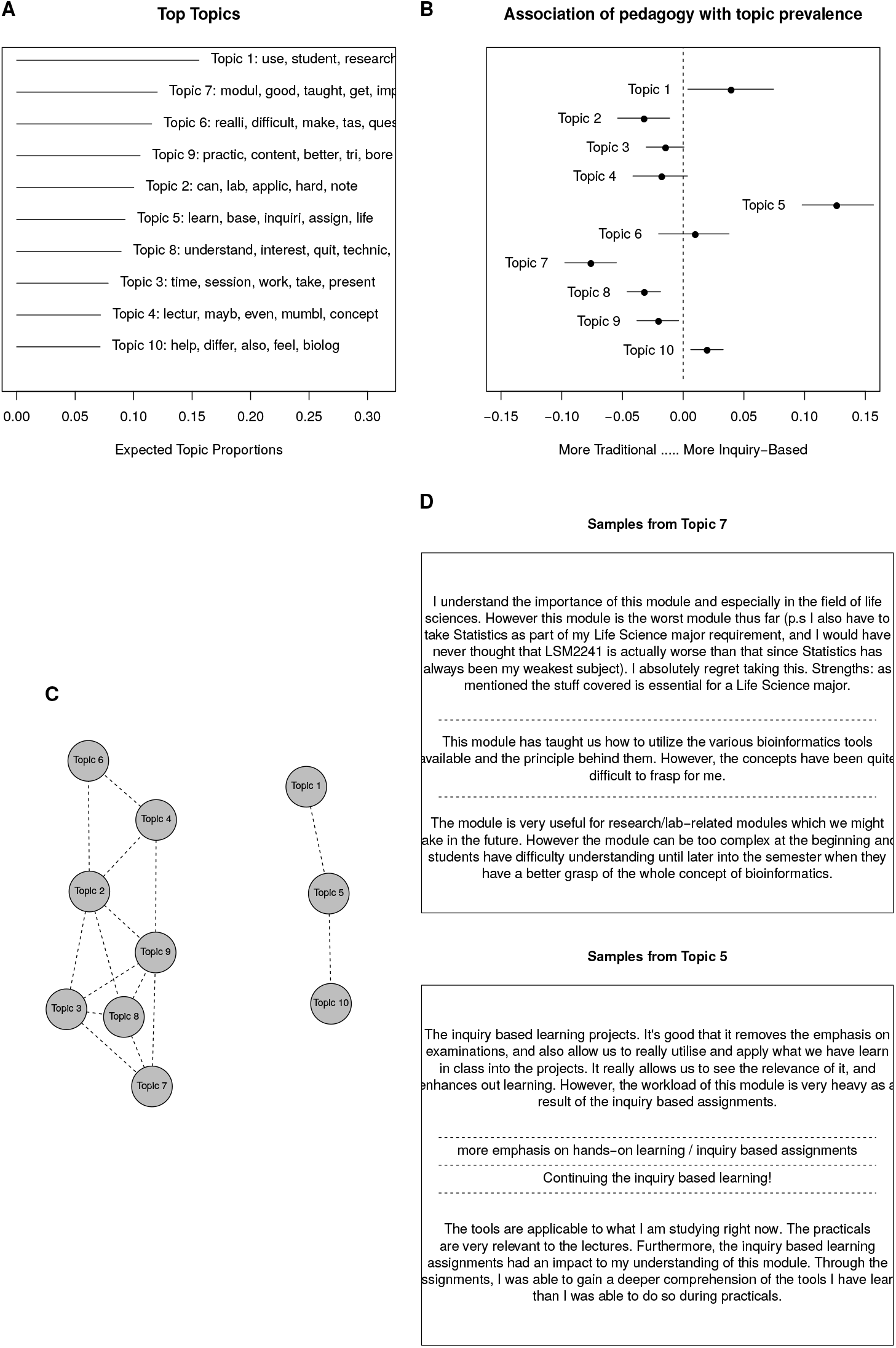
Structural topic modeling of student feedback. **A.** Topical prevalence for all topics, with sample stemmed words from each topic **B.** Relative prevalence of topics depending on pedagogy. Topics more strongly represented in the inquiry-based pedagogy are to the right, those more prevalent in the traditional pedagogy are to the left. **C.** Graph-based representation of correlation among topics. Highly correlated topics are connected; the three topics most associated with IBW are mutually correlated but distinct from the others. **D.** Sample comments from two topics most strongly associated with traditional (top) and inquiry-based (bottom) pedagogy in this module

### Observations of learning and development

When we designed the first assignment, we asked students to state a hypothesis for their inquiry. While students had received instruction on the nature of scientific hypotheses, they still found this part of the assignment extremely difficult and vague. We were hoping that students would use concepts (such as homology) from the lectures to frame their hypotheses. Students who formulated a hypothesis and then chose an appropriate test or tests might write passages such as: “We hypothesise that our given sequence is similar to annotated sequences in established databases and that we can therefore characterise our sequence by finding similar sequences using BLAST. Because it might be a partial cDNA sequence, we will initate our searches against protein databases”. While we received some well-formulated hypotheses, we received many hypothesis statements that were wild guesses (e.g., the sequence is human). The most striking common mistake was to form a hypothesis around the results (e.g., the sequence is from a specific family of transporter protein genes). We referred to this pattern as “the retrospective hypothesis”.

Common weaknesses in the first assignment were both rhetorical and methodological. A frequent pattern was for students to report all of their work, in the order it was performed, in the written assignment. If a student had not decided on a specific analysis strategy, he or she often performed analyses, and reported results, in the same order they were were covered in class. Such an approach is likely to use (and report) all the learned methods rather than deliberate and choose the most appropriate. This pattern is both rhetorically and scientifically weak, and it reflects a common misunderstanding possibly reinforced by a traditional laboratory course. Students are often taught to “let the data speak”, and admonished not to distort or manipulate their results. It is certainly essential for students of science to learn to recognise and avoid distortion, confirmation bias, *p* hacking, and the like. But data do not speak on their own, data need to be presented as results in the context of a sound scientific argument. Students should have good reasons for choosing the methods they employ, including reasonable hypothesis and expected results, and they should learn to justify their reasoning in their report writing.

The transformation of data from research into results of a research report is inherently rhetorical: it is the stuff of science. In a review session with students after the first assignment, we used the distinctions between a laboratory notebook and a research report to help the students reflect on this process. A laboratory notebook is chronological, while the narrative of a research study need not be. A good laboratory notebook documents all the work, while a published study may or may not. These distinctions may seem obvious to the reader, but need to be learned and internalised by students. An important part of this reflection was the importance of thoughtful study design, which is often aided by a vision of the research paper. Students who had considered the known properties of the sequence (it was presumed to be an incomplete cDNA) could often develop a productive strategy beforehand, starting with a blastx search of, say, the REFSEQ protein database. This both streamlined their work and strengthened their results, allowing their reports to more closely follow the chronological order of their efforts.

While this pedagogy used *writing* as the basis of student constructivism, research reports also contain figures and tables that need to be built. In the first report, many students showed the same weaknesses with tables and figures as they did with text, using (for example) multiple screenshots of BLAST searches rather than assembling results of multiple BLAST runs into concise figures or tables.

By the second assignment, most students had begun to internalise the need for essential rhetorical and scientific decisions. That learning was reflected in their writing: hypotheses were better formed, arguments were better made, and students made clear choices between analysis strategies rather than trying everything they had been taught. While students were evidently engaged with the inquiry-based work through the semester, their questions became more sophisticated and their group discussions more animated. In the second assignment, we noticed students making different choices based on what they had learned in the first assignment. Some students chose to focus on potential species of origin for their sequence, while other students developed phylogenetic trees of protein families.

In the third assignment, students worked in small groups to develop homology models of their sequences. Usually some sequences failed in this assignment to produce a good model. Groups handled this failure very differently, with some groups choosing to write only about the sequences that succeeded, while other groups organised their inquiry as a “compare and contrast” setting between the different sequences.

Our overall impression was that over time the students took increasing responsibility for the scientific decisions of their inquiry, and increasing ownership of the work product. We did not measure the effect of IBW pedagogy on student learning, so we cannot say whether the patterns we observed were a result of pedagogical change.

### Student feedback

Student Evaluations of Teaching (SETs) are a regular feature of university education. We decided to use SETs in this project specifically because SETs do not prompt students regarding the inquiry-based work, so provide a consistent forum for students to raise it of their own accord. SETs are problematic instruments when used to evaluate teacher performance, because open-ended feedback is inherently confounded with student biases (Boring, 2017; Boring, Ottoboni, & Stark, 2016). Here, however, the Structural Topic Modeling methodology allowed us to systematically identify topics from open ended survey questions that were associated with the pedagogical innovation of inquiry-based writing. The clarity of these results suggests that structural topic modeling may be a powerful tool to examine SET data in other contexts, including examination of student biases and topics associated with teacher performance ratings.

In the 2013-2014 SET form, students were asked to distinguish the best aspects of the module from those most needing improvement. The topic most associated with inquiry was not only overwhelming from students rating the best aspects of the module, but often showed student reflection on how the inquiry-based writing helped them think deeply and more clearly about the subject matter of the module.

### Objections and weaknesses

The Moskovitz & Kellogg (2011) proposal for IBW in the laboratory course has led to objections from scientific and writing educators. Michael Goggin (a physicist) argued that the double-blind assessment would undermine the teaching of science (see Goggin (2011) and response), and undervalues the writing involved in documentation. In our view this objection misunderstands the purpose for reforming laboratory writing to be more like authentic scientific inquiry, but does suggest a related concern: might the double-blind setting penalize students who performed correct work but wrote poorly, or reward students who wrote their way out of a poorly executed experiment? We did not examine that issue directly here, but are testing the correspondence between blinded and unblinded assessment in a separate study.

From the humanities, Catherine Prendergast has argued in support of Goggin that compositionists advocating for laboratory-centered Writing to Learn undervalue the importance of manual activities in the lab for constructive learning (Prendergast, 2013). She further notes that in her experience observing undergraduate students and mentors in a Research Experience for Undergraduates (REU) program, students learned a great deal by practicing manual laboratory activities, even though the REU required little writing. The REU program is similar to the Undergraduate Research Opportunities in Science (UROPS) program at NUS. As several of us are practicing research scientists, we appreciate the learning value of hands-on lab work, and the importance of the apprenticeship experience to develop “good hands”. But as Prendergast observes, the REU program (like the UROPS program at NUS) includes a regular schedule of constructive activities: presenting at group meetings, providing research updates, chalk talks, and perhaps journal clubs. The REU program requires a poster presentation instead of a written report; the NUS UROPS program requires a written report. The balance between different *forms* of composition expected in genuine research-based education will vary from program to program, and indeed from lab to lab.

Even given the exceptional value of research-based education such as REU and UROPS, students still benefit from laboratory classrooms. The laboratory classroom experience not only provides skills training that students can use when they take up research-based education, but also may help students decide whether to enter a research-based education program at all. A laboratory classroom that exposes students not just to the range of technical skills working scientists need to practice, but to reasoning about the open-ended possibilities of scientific inquiry, is a setting that we believe better prepares students for research-based education.

For pedagogical purposes, the original proposal suggested that students write not full lab reports but component parts to be dealt with separately depending on the level of student expertise. This suggestion dovetails with positions in the debate over inquiry-based learning that emphasize appropriate scaffolding for the problem at hand (Furtak, Seidel, Iverson, & Briggs, 2012; Hmelo-Silver, Golan Duncan, & Chinn, 2007; Zhang, 2016). For similar reasons, the difficulty of problems should be controlled so that appropriate problems are given to the students. While we did control the problem difficulty, we did not adopt the original proposal’s recommendation to limit writing to component parts of a paper; our decision not to do so increased stress for both students and teaching staff. The writing assignments were very difficult to mark in a timely manner, and because the lead instructor (GTK) had committed to reviewing and moderating marks for all students the workload was overwhelming.

In the third assignment we asked students to reflect as a group on the decisions they had made earlier as individuals. We wanted to assign students a reflective exercise, and the group inquiry seemed like an opportunity for it. In retrospect, we think this was a mistake for two reasons. First, that reflection doesn’t fit into the inquiry of the group assignment, and creates an unnatural rhetorical challenge in the assignment. In addition, some students were embarrassed about the choices they had made earlier, which led to tension in some groups. A separate reflection exercise outside of the IBW would have been a suitable alternative.

In the implementation described here, we encountered several additional issues. Student backgrounds varied widely, with a majority of the cohort majoring in life sciences (and having generally weaker mathematical training) and a minority of students majoring in statistics, computer science, and computational biology (with more quantitative skills but weaker life sciences training). The course did not require any prior programming experience and did not teach computer programming, but students with programming experience generally had a much easier time with the mathematical and algorithmic concepts of the class. Student heterogenity is a factor in any classroom, but we do not think we accounted for it sufficiently in designing the IBW implementation.

### Potential for IBW laboratories in other disciplines

Controlled variation of experimental problems can be created via computer simulations or virtual laboratories (Kuehn, 2018; Potkonjak et al., 2016), which have had a powerful impact on medical education, distance learning, and massive open online courses (MOOCs). A computer-based laboratory thus offers a possible setting for inquiry-based writing where multiple variations on a scenario can be developed without raising the safety concerns of the analogous experimental chemistry or physics laboratory. However, mimicking the problems of an experimental laboratory through computer simulations or virtual environments will lead to inauthentic student experiences. In an inherently computational subject such as bioinformatics, on the other hand, authentic problems can be computer-generated and solved in a laboratory classroom.

More broadly, any laboratory course devoted to experimental data *analysis* rather than experimental data *generation* could utilise the strategy we outline here to generate double-blinded individualised problem sets for students, assessed by inquiry-based writing assignments. Problem sets in biology could be generated by simulation (such as evolutionary games, simulated DNA sequencing data from genomics experiments) or by providing real experimental data available in excess (such as microscopy images).

Upper level classes also present opportunities for inquiry unavailable here. These courses did not require any prerequisite in computer-based analysis, and we did not at the time use Galaxy (Afgan et al., 2018); either would allow for more wide-ranging analysis as a basis for for inquiry-based writing assignment.

Our work here focuses on the laboratory classroom, as did Moskovitz & Kellogg (2011), but field-based courses can naturally provide individualised and blinded problem sets. Field-based courses may therefore provide opportunities for inquiry-based writing on experimentation without changing the experimental setting at all. While field-based learning may have inherent safety concerns, those concerns are not altered by the nature of assessment.

## Supporting information

Supplementary file 1: first assignment

Supplementary file 2: second assignment

Supplementary file 3: third assignment

## Acknowledgements

This work was supported by a Learning Innovation Fund – Technology grant (C-154-000-052-511) from the National University of Singapore to Greg Tucker-Kellogg

