## Supplementary file 1: first assignment for "Developing critical thinking in STEM education through inquiry-based writing in the laboratory classroom"

### LSM2241 CA1(a)
### Inquiry-based individual writing assignment: BLAST & Profiles

#### Background

You have each been given a DNA sequence which may code for a protein, in whole or in part, in any frame. No one, including the teaching staff, know what your sequence is, and no two students are given the same sequence. Through these inquiry-based assignments, you will characterize your sequences by employing various bioinformatics tools you have studied to this point. The same sequences, and the results of the first inquiry, will also be used for the later inquiry-based work in this module.

This inquiry gives you the opportunity to explore both your sequences and the tools of sequence analysis. You should plan your analysis based on any hypothesis or queries you might have about your sequences, conduct the analysis using appropriate bioinformatics tools, and interpret the results you obtain. Your assignment is a written report in PDF format. The written assignment is modeled on several characteristic parts of a scientific paper. You should report what you can infer about your sequence from your analysis, and why you are able to make claims about it. Many things can be discussed in your paper. What hypothesis did you start with? What was your analysis strategy? How did you interpret the results? Why did you use the results of some tools but not others? *Ultimately, what do your results tell you about the identity and characteristics of your sequence, and what else would you explore further?*

For this assignment, your analysis should be built around the BLAST suite of programs (covered in L05 and P05) and profile-based approaches (L06 and P06), along with other tools you have been taught to this point. You may also choose to use tools we *haven't* covered, within the general area of sequence database search and sequence profiles. Subject your sequences (DNA or protein sequences) to analysis using BLAST to get more information about your sequence and its relation to other sequences. Employ profile search tools such as PSI-BLAST or other profile-based methods to identify distant relatives and detect domain structure in your sequence. Consider how a profile search may allow you to characterise your sequence better than you were able with ordinary BLAST programs.

Here are some suggestions of things you might explore. Although the list below may appear to emphasise the tools, you should be considering the tools as means to an end: interpretable results! You have no obligation to explore all of the items listed below, or for that matter *any* of them. This is to just provide some ideas.

- Explore different flavours of BLAST and the consequence of your choice on results.
- Does choosing a different search database make a difference? When would you choose a different database?

- Did you use default BLAST parameters? If you did not, what modification did you make and how did it affect the interpretation of results?
- If you choose to use a tool like PROSITE-SCAN, what sort of choices might you make about the sequence or sequences to submit?
- Make a multiple sequence alignment from some of your Blast hits. Create a PSSM from it using EMBOSS PROPHECY.
- Test your sequence, or a multiple sequence alignment, using HMMER.
- If you created a PSSM, visualise it using the tools at the NCBI.
- Compare the PSSM you created with any PSSM in the CDD that were found in your original blast.
- Score your sequence against the PSSM you made from blast results of your sequence. Also score it against a PSSM from the CDD, if one was identified in your original BLAST. How do they compare?

#### Assignment

For the assignment, characterize your sequence using bioinformatics tools you have studied so far. Write a short report describing and sufficiently elaborating on your approach, your analysis, your results, and your interpretation. The report may contain any of the usual sections of a scientific paper (Abstract, Introduction, Methods and Materials, Results, and Discussion), but you may wish to skip or dramatically shorten some sections: the marking (described below) will emphasise the Methods & Materials and Results sections.

As a reminder, you are bound by the NUS Honour Code. Copying answers is pointless, because you each have a different problem. Copying text or figures without attribution is plagarism. If you have any questions about what constitutes plagarism, please review the first lecture materials and ask the teaching staff.

#### Submission of Assignment

Your paper should be no more than **four A4 pages** (**12 point font**), excluding any figures or tables you decide to include.

Please submit the assignment as a **PDF document**, with the title in the format of "CA1a-*YourMatricNumber*" (e.g. "CA1a-A0123456.pdf") to the upload folder.

Your report must be received by **11:59 PM on 2 October 2013 (Wednesday).**

### Assessment

**The assignment will be assessed according to the criteria below. Expectations of work marked "excellent" are provided.**

**Rationale -** An excellent paper will provide a clear hypothesis and a well explained rationale for why a search strategy was chosen. An excellent paper will explain and defend choices of tools used.

**Implementation -** An excellent paper will describe how the strategy was implemented. The implementation will use the tools correctly. Input parameters will be used appropriately, and significance will be evaluated without prejudice. If the strategy is described with the aid of a flow chart or figure, an excellent paper will support that flow chart or figure in the text.

**Interpretation -** An excellent paper will draw well defended conclusions about the sequences (domains, relatives, families), not hesitating to make supportable statements, but clearly distinguishing speculation from well supported inference.

**Organization:** An excellent paper will provide a well structured and comprehensive argument. The work will be organized into paragraphs that support the structure of the argument. Figures or tables will be clearly presented, with informative legends.

**References:** An excellent paper will provide a useful and correctly formatted bibliography for the reader. All use of figures, data, or text will be properly attributed and cited. No figures will be used without permission.

**Grammar, usage, and punctuation:** An excellent paper will use proper English, correct grammar and complete sentences.

| | **Nonexistent** | **Poor** | **Fair** | **Good** | **Very Good** | **Excellent** |
|---|---|---|---|---|---|---|
| Rationale | 0 | 4 | 8 | 12 | 16 | 20 |
| Implementation | 0 | 4 | 8 | 12 | 16 | 20 |
| Interpretation | 0 | 4 | 8 | 12 | 16 | 20 |
| Organization | 0 | 4 | 8 | 12 | 16 | 20 |
| References | 0 | 2 | 4 | 6 | 8 | 10 |
| Grammar, usage, and punctuation | 0 | 2 | 4 | 6 | 8 | 10 |
| **Total** | **0** | **20** | **40** | **60** | **80** | **100** |

**Late reports** will be penalized after marking according to the following table. The time of submission will be determined by the timestamp on the file in IVLE. Please be sure you know how to convert files to PDF in order to submit them to the system.

| **Late by** | **Penalty (points deducted)** |
|---|---|
| ≤ 6 hours | 10 |
| 6 hours < *timestamp* ≤ 12 hours | 25 |
| 12 hours < *timestamp* ≤ 24 hours | 50 |
| 24 hours < *timestamp* ≤ 48 hours | 75 |
| > 48 hours | 100 |
