## Supplementary file 2: second assignment for "Developing critical thinking in STEM education through inquiry-based writing in the laboratory classroom"

### LSM2241 CA1(b)
### Inquiry-based individual writing assignment: Molecular Evolution and Phylogenetics

#### Assignment

The work you have done to date may have given you a sense of whether your sequence belongs to a family of sequences, and what the characteristics of that family are.

A common next step with multiple sequence alignment and profiles is to develop a phylogenetic tree to explore evolutionary relationships. For the assignment, “your sequence” can refer to the DNA sequence provided or (with justification provided) a protein sequence based on your earlier work.

1. Gather (via BLAST, PSI-BLAST, or the profile method you used earlier) a set of sequences related to your sequence. Choose sequences that you think might make for an interesting evolutionary comparison.
2. Create and edit a multiple sequence alignment of those sequences, including your sequence. You may choose to focus on a particular region of sequence, so long as your provide a well-founded rationale for your choice.
3. Use MEGA or another phylogenetic tool to infer a phylogeny from the sequences.
4. Interpret, to the extent possible, where your sequence fits in the tree you have just developed.

For the assignment, write a short paper (corresponding to the “Methods and Materials” and “Results” section of a scientific article) describing your hypothesis, approach, and interpretation of your results (what you have found out about your sequences).

#### Submission of Assignment

Your paper should be no more than **two A4 pages** (**12 point font**), excluding references, and any figures or tables you decide to include. The total number of figures and tables must not exceed three (that is, if there are two figures, there can be only one table, and if there are three figures, there can be no tables).

Please submit the assignment as a **PDF document**, with the title in the format of “CA1b-*YourMatricNumber”* (e.g. “CA1b-A0123456A.pdf”) to the upload folder.

Your report must be received by **11:59 PM on 23 October 2013 (Wednesday).**

### Assessment

#### The assignment will be assessed according to the criteria below. Expectations of work marked “excellent” are provided.

**Rationale -** An excellent paper will provide a clear hypothesis and a well explained rationale for the approach was chosen. An excellent paper will explain and defend choices of tools used.

**Implementation** - An excellent paper will describe how the strategy was implemented. The implementation will use the tools correctly. Input parameters will be used appropriately, and significance will be evaluated without prejudice. If the strategy is described with the aid of a flow chart or figure, an excellent paper will support that flow chart or figure in the text.

**Interpretation** - An excellent paper will draw well defended conclusions about the phlygoenty, not hesitating to make supportable statements, but clearly distinguishing speculation from well supported inference.

**Organization:** An excellent paper will provide a well structured and comprehensive argument. The work will be organized into paragraphs that support the structure of the argument. Figures or tables will be clearly presented, with informative legends. The paper will be submitted in the required format.

**References:** An excellent paper will provide a useful and correctly formatted bibliography for the reader. All use of figures, data, or text will be properly attributed and cited. No figures will be used without permission.

**Grammar, usage, punctuation, and style:** An excellent paper will use proper English, correct grammar and complete sentences. The writing will be clear, consise, and coherent.

| | **Nonexisten t** | **Poor** | **Fair** | **Good** | **Very Good** | **Excellent** |
|---|---|---|---|---|---|---|
| Rationale | 0 | 4 | 8 | 12 | 16 | 20 |
| Implementation | 0 | 4 | 8 | 12 | 16 | 20 |
| Interpretation | 0 | 4 | 8 | 12 | 16 | 20 |
| Organization | 0 | 4 | 8 | 12 | 16 | 20 |
| References | 0 | 2 | 4 | 6 | 8 | 10 |
| Grammar, usage, | 0 | 2 | 4 | 6 | 8 | 10 |

| | | | | | | |
|---|---|---|---|---|---|---|
| and punctuation | | | | | | |
| **Total** | **0** | **20** | **40** | **60** | **80** | **100** |

**Late reports** will be penalized after marking according to the following table. The time of submission will be determined by the timestamp on the file in IVLE.  Please be sure you know how to convert files to PDF in order to submit them to the system.

| **Late by** | **Penalty (points deducted)** |
|---|---|
| ≤ 6 hours | 10 |
| 6 hours < *timestamp* ≤ 12 hours | 25 |
| 12 hours < *timestamp* ≤ 24 hours | 50 |
| 24 hours < *timestamp* ≤ 48 hours | 75 |
| > 48 hours | 100 |
