## Supplementary file 3: third assignment for "Developing critical thinking in STEM education through inquiry-based writing in the laboratory classroom"

### LSM2241 CA1c
### Inquiry-based group writing assignment:
### From Sequence to 3D Structure!

For this final assignment you will work in your project group (5-6 students per group) as registered in IVLE.

#### Background

Some of you will probably have worked on sequences that gave good profiles and strong similarity to well annotated sequences. Others probably did not. This is perfectly OK. In fact, it's how science works: sometimes the data are interpretable, other times not. Your task in this assignment is twofold: critical evaluation of individual profile and phylogenetic analyses, and homology modeling to see if any of your sequences can be understood in 3D structure.

You are doing this in groups for two reasons. First, by working in teams everyone should have a good opportunity to develop a homology model of at least one sequence. Second, by working together, you may be able to understand how the structural modelling relates to your previous work. You should compare and contrast the previous work you did on these sequences to see if you can identify and "best practices" that you can take into your professional lives and other studies.

#### Assignment

All of your individual work should be considered to inform this project. It will help you in formulating your hypothesis. You should each read the CA1a and CA1b reports from the other team members, and critically assess what factors (either of the sequences or the choices you made in analysis) led to deeper insight into the sequences.

For the homology modelling,

1. Using the results of your previous work, and any concepts you have learned about homology modelling, choose sequences (or subsequences) to attempt homology models.
2. Use SWISS-MODEL, MODELLER, or some other homology modelling tool to attempt homology modelling of these sequences.
3. Based on the results you obtain, refine your sequences, resubmit, and refine your results.

For the assignment, write a short paper (corresponding to the "Methods and Materials" "Results", and "Discussion" sections of a scientific article) discussing your previous work, describing your hypothesis, approach, and interpretation of your results.

### Submission of Assignment

Your paper should be no more than **five A4 pages** (**12 point font**), excluding references, and any figures or tables you decide to include. The total number of figures and tables must not exceed four (that is, if there are two figures, there can be only two tables). These limits include any appendix.

Please submit the assignment as a **PDF document**, with the title in the format of "CA1c-*YourProjectGroup"* (e.g. "`CA1c-C01.pdf`") to the upload folder.

Your report must be received by **11:59 PM on 8 November 2013 (Friday).**

### Assessment

The assignment will be assessed according to the criteria below. Expectations of work marked "excellent" are provided.

**Critical reasoning** - An excellent report will show evidence of critical thinking about the work you have done over the semester. Critical thinking in this case means "reasonable reflective thinking focused on deciding what to believe or do" See http://faculty.education.illinois.edu/rhennis/SSConcCTApr3.html for a set of useful bullet points.

**Implementation** - An excellent paper will use correctly the tools you have studied, with justified parameter selection, hypothesis formulation, and interpretation of results. We encourage you to think about your initial results and consider how you might refine your approach. Refinements based on initial results are both a form of implementation and (can be) a form of critical reasoning.

**Evaluation** - An excellent paper will draw well defended conclusions about the constructed model, not hesitating to make supportable statements, but clearly distinguishing speculation from well supported inference. You should also evaluate the model critically. *A good argument that a sequence cannot be modeled successfully is better than a bad justification for a weak model*.

**Teamwork:** An excellent report will demonstrate clear roles for different members of the team and a demonstration of contributions by each member of the team. You should include a brief section at the end with the title "Authors' contributions", with each team member's contributions to the work. You will be assessed *as a team;* the contributions section is not to assess individuals but to understand how you worked together. For example, one person might be responsible for organizing the report draft, another for running SWISS-MODEL, one for running PROCHECK or MOLPROBITY, one for organising the compare/contrast section, etc.

**Organization:** An excellent report will move smoothly through the subjects discussed, covering sufficient detail without repeating the material of the lectures or practicals.

**References:** An excellent paper will provide a useful and correctly formatted bibliography for the reader. All use of figures, data, or text will be properly attributed and cited. No figures will be used without permission.

**Grammar, usage, punctuation, and style:** An excellent paper will use proper English, correct grammar and complete sentences. The writing will be clear, concise, and coherent.

| | **Nonexistent** | **Poor** | **Fair** | **Good** | **Very Good** | **Excellent** |
|---|---|---|---|---|---|---|
| Critical Reasoning | 0 | 4 | 8 | 12 | 16 | 20 |
| Implementation | 0 | 4 | 8 | 12 | 16 | 20 |
| Evaluation | 0 | 4 | 8 | 12 | 16 | 20 |
| Teamwork | 0 | 2 | 4 | 6 | 8 | 10 |
| Organisation | 0 | 2 | 4 | 6 | 8 | 10 |
| References | 0 | 2 | 4 | 6 | 8 | 10 |
| Grammar and usage | 0 | 2 | 4 | 6 | 8 | 10 |
| **Total** | **0** | **20** | **40** | **60** | **80** | **100** |

**Late reports** will be penalized after marking according to the following table. The time of submission will be determined by the timestamp on the file in IVLE. Please be sure you know how to convert files to PDF in order to submit them to the system.

| **Late by** | **Penalty (points deducted)** |
|---|---|
| ≤ 6 hours | 10 |
| 6 hours < *timestamp* ≤ 12 hours | 25 |
| 12 hours < *timestamp* ≤ 24 hours | 50 |
| 24 hours < *timestamp* ≤ 48 hours | 75 |

| | |
|---|---|
| > 48 hours | 100 |
